# Characterization of a thalamic nucleus mediating habenula responses to change in illumination

**DOI:** 10.1101/047936

**Authors:** Ruey-Kuang Cheng, Seetha Krishnan, Qian Lin, Caroline Kibat, Suresh Jesuthasan

**Author notes:** equal contribution (Ruey-Kuang Cheng, Seetha Krishnan).

## Abstract

**Background:** Neural activity in the vertebrate habenula is affected by changes in ambient illumination. The nucleus that links photoreceptors with the habenula is not well characterized. Here, we describe the location, inputs and potential function of this nucleus in larval zebrafish.

**Results:** High-speed calcium imaging shows that onset and offset of light evokes a rapid response in the dorsal left neuropil of the habenula, indicating preferential targeting of this neuropil by afferents mediating response to change in irradiance. Injection of a lipophilic dye into this neuropil led to bilateral labeling of a nucleus in the anterior thalamus that responds to onset and offset of light, and that receives innervation from the retina and pineal organ. Lesioning the neuropil of this thalamic nucleus reduced the habenula response to light. Optogenetic stimulation of the thalamus with channelrhodopsin-2 caused depolarization in the habenula, while manipulation with anion channelrhodopsins inhibited habenula response to light and disrupted climbing and diving that is evoked by irradiance change.

**Conclusions:** A nucleus in the anterior thalamus of larval zebrafish innervates the dorsal left habenula. This nucleus receives input from the retina and pineal, responds to increase and decrease in irradiance, enables habenula responses to change in irradiance, and may function in light-evoked vertical migration.

## Background

The habenula is an evolutionarily conserved structure [1] that influences multiple behaviors, ranging from fear [2–4], to learning [5–7], addiction [8], sleep [9], aggression [10,11] and performance under stress [12]. One function of the habenula is to regulate the release of broadly-acting neuromodulators such as serotonin, dopamine, epinephrine and histamine [12–15]. To precisely control these neuromodulators, the habenula integrates diverse variables including internal state, reward value and sensory stimuli. This information reaches the habenula from distinct sources. For example, circadian time is transmitted to the habenula by hypocretin-secreting neurons located in the hypothalamus [16]. Negative reward or punishment is conveyed by neurons of the entopeduncular nucleus (internal segment of the globus pallidus) [17]. Olfactory stimuli evoke activity in the habenula [18,19] via a direct innervation of mitral cells from the olfactory bulb [20]. Light as well as loss of light can also cause activity in the habenula, as has been demonstrated in rat [21], mouse [22], pigeon [23] and zebrafish [18,24], but the neurons regulating habenula responses to changes in ambient illumination are not well defined.

The habenula is divided into two major regions based on pattern of connectivity. In mammals, these are called the medial and lateral habenula, while in fish these are the dorsal and ventral habenula [25]. In larval zebrafish, short pulses of red light cause asymmetric depolarization of the dorsal habenula, with a stronger response on the left side [18]. This response is lost in fish lacking eyes [18]. However, no direct pathway from the retina to the habenula has been documented [26,27]. By retrograde tracing in adult zebrafish, Turner et al proposed that the habenula receives input from the nucleus rostrolateralis [28], a diencephalic structure with retinal input that has been described in several ray-finned fish [29,30]. Because injections into both left and right habenula led to labeling of this structure, and also because of potential artifacts in labeling, the authors concluded that the source of light-evoked activity in the habenula could not be determined [28]. Here, we set out to characterize the nucleus by which ambient light affects activity in the habenula and to explore the function of this nucleus in innate responses to change in irradiance.

## Results

### Habenula afferents mediating response to change in irradiance target t he dorsal left neuropil

The zebrafish habenula consists of neurons surrounding neuropils that are innervated by afferent neurons located in different regions of the brain [20,25,28,31]. To identify neurons that mediate light-evoked activity in the habenula, we sought to determine which neuropils are affected by this stimulus; blue light was used as the stimulus, as the habenula has a strong response to this wavelength [32]. We first used wide-field microscopy, which has a large depth-of field and thus provides an overview of stimulus-evoked fluorescence. Imaging was carried out on *elavl3:GCaMP6f* fish, which have broad expression of the calcium indicator. The exposure time (5 msec; 200 Hz) was shorter than the rise time of GCaMP6f (~80 msec; see supplementary table 1 in [33]), so the initial image primarily reflected activity prior to effects of the stimulating light. With this approach, a rapid increase in fluorescence was detected in a discrete region in the dorsal left habenula following onset of the blue light used to excite the reporter (Fig. 1a-c; n = 5 fish). This suggests that onset of light activates a neuropil in the left habenula.

Two-photon microscopy was used next, as this allows higher spatial resolution imaging before, during and after delivery of a more precisely timed light stimulus. Imaging was carried out using *GAL4s1011t, UAS:GCaMP6s* fish, in which expression of the calcium reporter is restricted mainly to habenula neurons [19]. In agreement with wide-field microscopy, onset of light was found to trigger a response in the dorsal left neuropil of the habenula (Fig. 1d-h; blue pixels). Responses in the neuropil, which contains dendrites of habenula neurons, correlated with but preceded the response of the cell body of habenula neurons (Fig. 1i-k). These observations suggest that neurons mediating the habenula response to light reside outside the habenula and target the dorsal left neuropil.

In addition to responses to the presence of light, we also detected a response to the offset of light in the dorsal left neuropil (Fig. 1g, h; magenta pixels). As with the response to light ON, the light OFF response in the neuropil preceded the response in habenula neurons (Fig. 1k). Two different classes of response to light OFF could be seen in habenula neurons (Fig. 2). In one, the activity was suppressed during light ON and increased after offset of light; in the other, there was an increase in activity after the pulse of light but there was no decrease during the pulse (Fig. 2a-c). The former class is referred to as INH (for “inhibited”). All three classes of neurons – ON, OFF and INH – were more numerous in the dorsal left habenula as compared with the right (Fig. 2d), similar to what has been reported before for neurons that respond to the onset of red light [18]. Responses were seen in all ages examined (5 – 10 dpf) (Fig. 2e), indicating that this phenomenon is not restricted to early stages of nervous system development. These observations suggest that the dorsal left habenula is innervated by neurons that respond to light and darkness.

### A thalamic nucleus innervates the habenula

To identify afferents to the dorsal left neuropil, the lipophilic tracer DiD was injected here (Fig. 3a; n = 10 fish). This led to bilateral labeling of a cluster of neurons posterior to the zona limitans thalamica (ZLI) (Fig. 3b, c, Movie 1 (Additional File 1)). In contrast, when the right dorsal habenula was injected with DiD, label was seen primarily in the ipsilateral entopeduncular nucleus (Fig. 3d, Movie 2 (Additional File 2); n = 4 fish). We hypothesized that neurons innervating the dorsal left habenula belong to a thalamic nucleus, given their position relative to the ZLI, which marks the anterior limit of the thalamus. A feature of the thalamus in zebrafish is the presence of GABAergic neurons at the rostral margin [34]; in general, first-order thalamic nuclei contain GABAergic interneurons that synapse onto the axons of incoming sensory neurons [35]. Thus, if the habenula-projecting nucleus were a thalamic nucleus, it would be expected to contain GABAergic neurons that extend neurites into its neuropil. Consistent with this, DsRed that was driven by the *gad1b* promoter [36] was seen in cells posterior to the ZLI and in the neuropil of the anterior thalamus (Fig. 3e; Movie 3 (additional file 3)). Additionally, immunofluorescence with the GAD65/67 antibody labeled the neuropil containing retrogradely labeled habenula afferents (Fig. 3f). Given the location and the presence of GABAergic neurons, the neurons that innervates the dorsal left habenula appear to be located within an anterior thalamic nucleus.

Thalamic nuclei usually contain glutamatergic projection neurons [35], but may in rare cases extend GABAergic projections [37]. When DiD was injected into the dorsal left habenula, approximately 45% of retrogradely labelled thalamic neurons expressed eGFP under the control of the vGlut2 GAL4 driver [36] (arrowheads, Fig. 3b; Movie 4; n = 3 fish), consistent with projection neurons being glutamatergic. We asked whether any of the afferent neurons might be GABAergic, as this is one possible mechanism for the suppression of activity seen in INH neurons. However, no retrograde DiD label was seen in thalamic cells labeled by *gad1b:DsRed* (see Movie 1 and Movie 4), nor were there DsRed labeled neurites in the dorsal left habenula neuropil (see Movie 3), as would be expected if there was innervation by GABAergic neurons. Moreover, when the fixable dye CM-DiI was injected into the dorsal left habenula, followed by immunofluorescence with the antibody to GAD65/67, no double-labeled cells were seen (Fig. 3f, n = 6 fish). It is unclear whether this is due to the low probability of labeling the relevant GABAergic cells with lipophilic tracing or with the transgene, or, more simply, if there are no GABAergic projections from the thalamus to the habenula. It is evident, however, that the anterior thalamus sends glutamatergic projections to the habenula.

### Irradiance change evokes activity in the anterior thalamus

If the anterior thalamic nucleus mediates illumination-dependent activity in the habenula, neurons here should respond to increase and decrease of illumination. To test this, calcium imaging was carried out in *GAL4s1020t, UAS:GCaMP6s* transgenic fish, which expresses the calcium indicator in thalamic neurons. Given the depth spanned by thalamic neurons innervating the dorsal left habenula (Fig. 3c), multiple planes were imaged, using a piezo-drive for fast focusing and resonant scanning for fast acquisition. A response to increase or decrease in illumination was detected in cell bodies of the anterior thalamus (Fig. 4a-j) in all fish imaged (n = 10). A minority of cells (yellow) responded to both increase and decrease. Responses could also be seen in the thalamic neuropil (Fig. 4k-m), which would be expected to receive driver (i.e. sensory [35]) inputs. The response to light was stronger for blue than for red light (Fig. 4n-r) as was seen previously with lower resolution whole-brain imaging [32]. Thus, calcium imaging supports the hypothesis that the anterior thalamic nucleus mediates responses to both increase and decrease in illumination, and also shows that it is more responsive to blue rather than red light.

The habenula response to light has been reported to be eye dependent [19]. To test if the habenula-projecting thalamic nucleus is innervated by retinal ganglion cell (RGC) axons, DiI was injected into the retina in fish where DiD had been injected into the dorsal left habenula. RGC axons could be seen to intermingle with neurites from DiD labeled neurons in the neuropil of the anterior thalamus (Fig. 5a, Movie 5 (Additional File 5)). This terminal field may include the previously described AF4 and AF2 [26,27], given the position anterior and medial to the optic tract. Consistent with this, two regions within the anterior thalamus neuropil of *elavl3:GCaMP6f* fish (in which retinal ganglion cells are labelled) responded to change in illumination, with increase in irradiance causing activity more dorso-caudally while decrease caused activity more rostro-ventrally (Fig. 5b-h).

If the anterior thalamic nucleus functions as a relay for information from the retina to the habenula, the thalamic response to light should be absent in fish lacking eyes. Surprisingly, although the response to light ON was reduced, a response to light OFF could still be detected (Fig. 6a-l). This implies that there could be non-retinal inputs to this nucleus. One potential source may be the pineal organ, as injection of DiI into the pineal led to labelling of axons that extended into the neuropil of the anterior thalamus (Fig. 6m, n; Movie 6 (Additional File 6)), and the pineal has a response to light OFF (Fig. 6o, p). An OFF response was seen in the habenula in fish lacking eyes (Fig. 6q-s), consistent with the habenula being a target of the anterior thalamus, and the retina not being the sole source of sensory input to this nucleus.

### Lesion of the anterior thalamic neuropil inhibits habenula response to illumination change

To test if the anterior thalamic nucleus contributes to light-evoked activity in the habenula, we lesioned the anterior thalamic neuropil with a femtosecond laser. *elavl3:GCamp6f* fish were used, to enable imaging prior to lesioning so that the thalamic neuropil could be visualized and the plane with the response to light identified. Repeated pulsing with the laser led to the formation of a cavitation bubble in the neuropil (Fig. 7a), a characteristic feature of two-photon lesioning of tissue [38,39], and resulted in a reduction of both ON and OFF responses in the habenula (Fig. 7b-g). As a control for specificity of the lesioning technique, we targeted the parapineal, a light-sensitive organ that is located adjacent to the left habenula and directly innervates the dorsal left neuropil (Fig. S1a (Additional File 7)). This did not have any significant effect on habenula response to blue light or darkness (Fig. S1), indicating that the lesioning technique used here does not cause indiscriminate damage to surrounding tissue. Moreover, these observations suggest that the parapineal does not have an essential role in habenula response to illumination conditions, consistent with the findings of Dreosti et al [18], whereas the anterior thalamus is required.

### Optogenetic manipulation of the thalamus affects habenula response to irradiance change

Physically lesioning the anterior thalamic neuropil with the femtosecond pulsed laser is a difficult experiment, due to the presence of a blood vessel in the neuropil: ~80% of lesioned animals could not be used due to bursting of this vessel. As an alternative method of silencing the thalamus, we developed a transgenic line expressing the anion channel rhodopsin ACR1 from *Guillardia theta* [40]. This channel generates a chloride current in the presence of green or blue light, thus hyperpolarizing neurons. We expressed this channel in thalamic neurons under the control of the *GAL4s1020t* driver; expression in the anterior thalamus was confirmed by the presence of the YFP tag (Fig. 8a, b). This channel can be actuated in larval zebrafish by blue or green light at a power density of approximately 3 µW/mm^2^ (see accompanying manuscript) [41]. With this level of light, fish expressing GtACR1 showed a reduced response in the dorsal left neuropil to light ON compared to non-expressing siblings (Fig. 8c-j), consistent with input from the thalamus being required.

Strikingly, there was a stronger response to the offset of light in GtACR1-expressing fish, compared to non-expressing siblings (Fig. 8g-k). This may be the result of depolarization at the termination of light-gated hyperpolarization, as has been reported for other light activated chloride channels [42,43] and for GtACR1 (see accompanying manuscript). This finding implies that depolarization of thalamic neurons can drive habenula activity. To test this more directly, we examined the effect of optogenetic activation of the thalamus with Channelrhodopsin-2 (Fig. 9). This experiment was carried out in fish lacking eyes, to prevent a visual response. Short pulses of blue light reproducibly caused an increase in fluorescence of GCaMP6f in habenula neurons of fish with expression of ChR2 in the thalamus (Fig. 9b, f). No change in fluorescence was seen in the absence of blue light (Fig. 9c, d), indicating that the activity is due to the stimulus. Some response was seen in fish without ChR2 expression (Fig. 9e), suggesting that a component of the habenula response may be due to non-ocular sensors such as deep brain photoreceptors [44,45]. The larger response in fish with ChR2 expression (Fig. 9g-h), however, is consistent with the hypothesis that a thalamic nucleus regulates activity in the habenula of larval zebrafish.

### Optogenetic manipulation of the thalamus disrupts an innate behavioral response to irradiance change

Finally, we asked whether the anterior thalamic nucleus might be involved in an innate behavior that is responsive to change of light. We hypothesized that one such behavior may be light-evoked vertical migration [44]. Larval zebrafish normally move upwards to the surface of a water column in the presence of blue or green light, but move downwards when the lights are switched off [32]. We tested the effect of optically manipulating the thalamus using the anion channel rhodopsins, reasoning that the presence of these channels would disrupt normal light-controlled responses: if no difference was seen, then the hypothesis should be rejected.

Fish expressing GtACR1 or GtACR2 under the control of the *GAL4s1020t* driver behaved differently from siblings (Fig. 10a, b). Rather than swimming upwards in the light, GtACR-expressing fish were seen to move downwards in the light and to swim upwards in the dark. This is reflected by a reversal in the correlation between position in the tank and illumination status in GtACR1 or GtACR2-expressing fish, in contrast to non-expressing siblings (Fig. 10c, e). One potential reason for GtACR-expressing fish to swim upward at the offset of light could be that there was a lack of space to move downwards, given their starting position near the bottom of the tank. To test whether space was a constraint, we plotted the direction of initial movement at light OFF as a function of position (Fig. 10d). Although GtACR1 fish were predominantly located near the base of the tank, a number were located in the middle of the tank, i.e. between a relative depth of 0.25 and 0.75, where they would have space to move up or down. A comparison of the behavior of fish in this region, using multilevel analysis to rule out nesting effects caused by repeated measures on the same fish [45], indicated significant difference between GtACR1-expressing fish and siblings X^2^ = 6.8958, p = 0.0088, df = 1). We also investigated whether the lack of climbing could be due to an inability to swim in the presence of light, given that the *GAL4s1020t* line can drive effector genes in motor neurons [46]. As shown in Fig. 10g-h, both GtACR1 and GtACR2 expressing fish moved less than siblings in the presence of light, although there was not a complete cessation of movement. Thus, while some of the loss of climbing in the presence of light could be due to nonspecific effects, the overall result, including the downward movement in the light and climbing at light offset is consistent with the hypothesis that the anterior thalamic nucleus has a role in climbing behavior that is normally triggered by light.

## Discussion

Imaging with wide-field and two-photon microscopy demonstrates that the dorsal left neuropil of the zebrafish habenula is stimulated by change in light, consistent with previous reports of an asymmetric response in habenula neurons to a flash of light [18]. Lipophilic tracing demonstrates that this neuropil is asymmetrically innervated by a nucleus in the anterior region of both left and right thalamus. The anterior thalamic nucleus receives input from the retina and pineal, and responds to change in irradiance. Lesion of the anterior thalamic neuropil or optogenetic silencing of the thalamus inhibited light-evoked activity in the habenula, while optogenetic stimulation of the thalamus drove activity in the habenula. Thus, by optical recording, anatomical tracing, optical manipulation and lesion, our data suggests that an anterior thalamic nucleus mediates the habenula responses to irradiance change in larval zebrafish.

The thalamic nucleus that projects to the habenula can be functionally separated into two domains, based on the response to light – excitation to light OFF in the anterior-ventral regions and excitation to light ON more dorso-posteriorly. This neuropil contains two previously defined targets of retinal ganglion cells, AF2 and AF4 [27], that have this location. AF4 is innervated predominantly by M3 and M4 retinal ganglion cells, which extend their dendritic tree into the proximal layer of the inner plexiform layer and are considered ON neurons [27]. AF2 is innervated by B1 retinal ganglion cells that have dendrites in the distal layer [27], and these may account for the OFF responses in the thalamus and habenula. The pineal may also be responsible for a component of OFF responses: pineal cells appear to depolarize in darkness, and pineal fibers innervate the thalamic neuropil of larval zebrafish, as has been reported for adult zebrafish [47].

As in the anterior thalamus, a response to the loss of light was seen in the habenula. This has a number of implications. Firstly, this suggests that darkness itself may be a stimulus, in which case the level of activity in habenula neurons during darkness prior to a light stimulus cannot be taken to be a “ground” state. Such activity may include what has been termed spontaneous activity [48], but may additionally reflect the current state of the animal (i.e. the effects of being in the dark, which is aversive to larval zebrafish [49]). Secondly, the fact that there is more than one class of habenula response to darkness implies that there may be more than one mechanism involved. In particular, the suppression of activity in the presence of light in INH neurons implies that a part of the OFF response could involve active inhibition. As yet, there is no evidence that there is direct hyperpolarization of habenula neurons during light ON. However, inhibition need not occur in the habenula, but could occur in the thalamus, where there are GABAergic neurons that extend neurites into the thalamic neuropil. Inhibition of thalamic OFF neurons by thalamic ON neurons, for example, could lead to the observed pattern in habenula INH neurons.

The thalamic nucleus mediating activity in the habenula may represent the nucleus rostrolateralis, as proposed by Turner et al [28]. The nucleus rostrolateralis was initially described as a dorsal thalamic nucleus that receives retinal input [50]. However, it was more recently suggested that this nucleus is an extension of the habenula, due to apparent innervation of the interpeduncular nucleus (IPN) [30]. We find no evidence that the nucleus identified here has a direct connection to the IPN. The *GAL4s1020t, UAS:GCaMP6s* line which was used for calcium imaging the thalamic response, for example, does not label axons extending to the IPN. Moreover, the *GAL4s1011t* driver, which labels the habenula neurons and axons that extend to the IPN, does not label the nucleus with retinal input. It is thus unclear whether the nucleus identified here is different from the nucleus rostrolateralis described in the butterfly fish, or if there was a labeling artifact in the tracing experiment [30].

While this manuscript was in review, it was suggested that light-evoked activity in the habenula is driven by input from the thalamic eminence (EmT) [51], an “ambiguous thalamic structure” [34] that has been proposed to give rise to the glutamatergic bed nucleus stria medullaris (BNSM) [34,52] or the ventral entopeduncular nucleus, a homolog of the globus pallidus [28]. However, the nucleus characterized here is distinct from the ventral entopeduncular nucleus, which is located more anteriorly and ventrally [28]. It also contains GABAergic neurons, and is thus unlikely to be the BNSM. It is possible that the nucleus here is an additional derivative of the EmT, although this remains to be demonstrated with lineage tracing. Intriguingly, Zhang et al [51] show that the retinal inputs to AF4 express the melanopsin-related gene *opn4xa*, consistent with our finding that the thalamic response to light is stronger for blue light relative to red light, and another report that the habenula response is stronger for blue light [32]. In mammals, melanopsin expressing retinal ganglion cells target a number of thalamic structures, including the intergeniculate leaflet and the margin of the lateral habenula [53]. The latter region may correspond to the para-habenular termination zone, which is located in the anterodorsal thalamic nucleus [54]. Whether either of these regions is homologous to the zebrafish nucleus described here remains to be determined.

Neurons in the anterior thalamus have a prominent sustained response to blue light (see Fig. 4a-e and [32]), and may be involved in a behavior that is evoked by blue light, which is vertical migration. This response is disrupted by expression of anion channelrhodopsins in the anterior thalamus, suggesting that the behavior is not independent of the thalamus. A limitation of this experiment, however, is that the driver line used also causes expression of the channel in spinal motor neurons [46]. Silencing of these neurons may contribute to reduced ability of GrACR1 and GtACR2 fish to move upwards in the light. However, the offset of light, which causes activity in networks containing light-gated chloride channels (see Fig. 8) [43,55], led to upward movement. This is unlikely to be due only to rebound activation of motor neurons, as there is a choice of which direction to move. Instead, the upward movement at light offset is consistent with the hypothesis that activation of the thalamus may drive vertical migration.

A projection from the thalamus to the habenula may be evolutionarily conserved in vertebrates. Using retrograde tracing with horseradish peroxidase, a projection from the dorsal thalamus to the habenula has been reported in a lizard [56] and in the frog [57]. In humans and rabbits, a thalamo-habenula projection was proposed many years ago based on degeneration experiments [58,59], but evidence with modern tracing techniques is lacking. Hints of a projection can be seen in a tracing experiment performed in rats [60], but this remains to be confirmed. The mesoscale mouse connectome project [61] also suggests that such a projection may exist, but the large volume of label means that the possibility of label from neighboring regions cannot be excluded. In humans, resting state functional magnetic resonance imaging indicates that the habenula and thalamus are directly connected [62,63]. However, it remains to be determined whether this connection is direct. The findings in lower vertebrates suggest that it may be worthwhile revisiting efferent connectivity of the anterior thalamus in mammals and investigating if and how this mediates non-visual responses to light.

## Conclusions

A nucleus in anterior thalamus of zebrafish enables habenula responses to increase and decrease in ambient illumination. This nucleus is innervated by the retina and pineal. It may function in vertical migration triggered by light.

## Methods

Experiments were performed in accordance with guidelines issued by the Institutional Animal Care and Use Committee of the Biological Resource Centre at Biopolis, Singapore.

### Fish lines

Zebrafish (*Danio rerio*) lines used for this study were: *UAS:GCaMP6s^sq205^*, SqKR11Et [2], sqKR4Et [64], *GAL4s1011t* [65], *GAL4s1020t* [65], *UAS:GCaMP3^sq200^*, *elavl3:GCaMP6f^a12200^*, *UAS:ChR2-eYFP [66]*, gad1b:DsRed [36], vGlut2a:GAL4 [36], UAS:eGFP, UAS:GtACR1 [55], UAS:GtACR2 [55], elavl3:GCaMP6f (Wolf et al, in press) and AB wildtype.

### Imaging

Zebrafish larvae were anaesthetized in mivacurium and embedded in low-melting temperature agarose (1.2-2.0 % in E3; egg water: 5 mM NaCl, 0.17 mM KCl, 0.33 mM CaCl2, 0.33 mM MgSO4) in a glass-bottom dish (Mat Tek). They were imaged on a Nikon two-photon microscope (A1RMP), attached to a fixed stage upright microscope, using a 25x water immersion objective (NA = 1.1). The femtosecond laser (Coherent Vision II) was tuned to 920 nm for GCaMP imaging. Stacks were collected in resonant-scanning mode with a 525/50 nm bandpass emission filter and with 8x pixel averaging; single-plane images were collected in galvano-scanning mode with 2x pixel averaging.

Light stimuli were generated by 5 mm blue LEDs (458 nm peak emission). They were powered by a 5 V TTL signal from a control computer and synchronized with image capture using a National Instruments DAQ board, controlled by the Nikon Elements software. Light intensity at the sample was 0.13 mW/cm^2^.

For widefield microscopy, excitation was provided by LEDs (Cairn OptoLED) at 470 nm. Images were captured on a Zeiss Axio Examiner with a 20x water immersion objective, using a Flash4 camera (Hamamatsu) controlled by MetaMorph. After background subtraction, change in fluorescence was measured using MetaMorph.

### Image data analysis

#### Initial Data Preprocessing

Data was analysed using custom written codes in Python. Raw images obtained were first registered using cross-correlation to correct for any vertical/horizontal movement artifacts. Then, a median spatial filter of size 3 was applied to remove spatial noise. A darker region outside the fish was chosen as the background and subtracted from the image to remove any signal that did not arise from GCaMP fluorescence. Non linear trends in the data were detrended using polynomials of order 2-3.

#### Pixel based analysis in single fish

In order to look at the overall spatial distribution of responses, which included both neuropils and cells, we performed clustering via *K-means* using the Thunder platform [67]. Data here were normalized into Z-scores by subtracting the overall mean and dividing by the standard deviation of each pixel over time and smoothed with a rolling window. Since pixel based analysis are sensitive to noise, and neighboring pixels with the same response could have varying standard deviation (in case of cell segmentation, pixels forming an ROI are averaged to obtain its intensity value), z-scores that account for both mean and standard deviation were used. Clusters obtained using pixel-based k-means analysis also provided the basis for the type of responses we looked for in segmented neurons.

#### K-means

*K-means* clustering was performed to identify pixels with similar responses profiles. This algorithm classifies the pixels into clusters, where the number of clusters, k, is chosen by the user. The end results are k cluster centers and labelling of pixels that belong to each cluster. Given the uncertainty of the optimal cluster number, an iterative approach was used to separate pixels relating to evoked responses versus pixels that do not (here referred to as independent clusters). The number of clusters were chosen to reveal as many stimulus related clusters as possible, until there was little change in the number and types of stimulus related clusters and increase in independent clusters. In normal fish, clusters related to evoked activity were easy to obtain. Clusters that are stimulus-independent were removed from the spatial and temporal plots for clarity. Examples of such clusters are shown in Fig. S2 (Additional File 8). In all cases, *K-means* cluster centers showing evoked responses to light ON were colored in shades of blue and those showing responses to light OFF were colored in shades of red. Pixels belonging to the cluster were colored similarly and superimposed on an average image of the plane analysed. In different datasets (Fig. 1d-e, Fig. 4a-f, 4n-o and 5c-e), this analysis provided an optimal k of 6-10. The 2-4 clusters that did not correspond to evoked activity were not included while plotting.

#### Cell Segmentation

Cells were manually segmented in ImageJ. The average intensity of pixels within an ROI across time were saved for further analysis. ΔF/F_0_ of the temporal traces were calculated by subtracting and then dividing by the mean of the total fluorescence during a baseline period (usually 10 seconds before first stimulus). A rolling window average was performed to smooth traces.

#### Classifying responses

Pixel based *K-means* analysis revealed many categories of responses to changes in irradiance. Using that as a basis, temporal traces from the cells were first broadly classified as those responding to light ON or light OFF. This was done by calculating their correlation coefficient to a square wave that is 1 when the light is ON or when light is OFF and 0 during other time periods. High correlation to these traces indicated that the pixel or cell is responding to light ON or OFF respectively (from multiple runs, a correlation coefficient of 0.4 and above seemed to provide accurate classification). Inspecting the cell traces in the ON and OFF categories revealed further classifications that could be made based on time of response (transient or sustained) and direction of response (excitatory or inhibitory). Cells responding transiently to both ON and OFF were also found. The temporal traces from the many categories in individual fish are plotted as heatmaps (for example, Fig. 2a, 4g). In experiments looking for the presence or absence of activity (effects of anterior thalamus neuropil ablation, parapineal ablation, enucleation, red vs blue response), the broad categories of ON and OFF were used. Spatial distribution of these categories are also plotted (for example, Fig. 2b, 4h).

#### Neuropil responses

Similar to the cell responses, pixels from habenula, thalamic neuropil and the pineal were similarly classified. Pixels from multiple fish were overlaid on each other and image transparency was adjusted to view the compiled response. Since locations of responses were largely similar and different classes spatially distinct in the neuropil of individual fish, the overlay did not mask any response.

#### Boxplots

Where possible, boxplots were plotted to show the full distribution of the data. The box in the boxplot ranges from the first quartile to the third quartile, and the box shows the interquartile range (IQR). The line across the box is the median of the data. The whiskers extend to 1.5*IQR on either side of the box. Anything above this range are defined as outliers and plotted as black diamonds in the plots.

#### % of active cells/pixels vs % of cells/pixels

% of active cell/pixels were calculated by dividing the number of active cells / neuropil pixels by the total number of cells / neuropil pixels. These provide an indication of the response across individual animals and have been shown as boxplots or individual data points. Histograms, on the other hand, display % of cells/pixels, which are obtained by dividing number of cells / neuropil pixels with a particular ΔF/F_0_ by the total number of cells / neuropil pixels.

#### Statistics

The Kolmogorov-Smirnov test (KS-test) was used to calculate the differences in distribution of amplitude or response duration. Histograms are shown in all cases. For non parametric paired distributions of number of cells, a Wilcoxon signed rank test was used and a Mann Whitney U test was used for independent data. Test statistic and p-values are reported.

### Neural tracing

DiD (Thermo Fisher Scientific) was dissolved in 50 µl ethanol to make a saturated solution. This was heated to 55°C for 5 minutes prior to injection into the fish that had been fixed in 4% paraformaldehyde. Fish were mounted in 2% low melting temperature agarose dissolved in PBS. The dye was pressure injected into the habenula under a compound microscope (Leica DM LFS), using a 20X water immersion objective. For labeling the retina, a saturated solution of DiI (Thermo Fisher Scientific) in chloroform was used. Injections were carried out under a stereomicroscope (Zeiss Stemi 2000). After injections, fish were stored at 4°C overnight to allow tracing, and then imaged with a 40x water immersion objective on a Zeiss LSM 710 confocal microscope.

CM-DiI (Thermo Fisher Scientific) was dissolved in ethanol (1 µg/µl). Fish were mounted in 2% agarose in E3, injected on a compound microscope, then allowed to recover in E3 at 28°C for 4 hours.

### Antibody label

Larvae were fixed in 4% para-formaldehyde/PBS overnight at 4°C. They were then rinsed in PBS. The brains were dissected out, and permeabilized using 1% BSA (fraction V; Sigma), 0.1% DMSO and 0.1% Triton X-100. The anti-GAD65/67 (Abcam ab11070, RRID:AB_297722; 1:500) has previously been used in zebrafish [2,68]. The brains were incubated in the primary antibody overnight, rinsed several times in PBS, then incubated in secondary antibody (Alexa 488 goat anti-rabbit; 1:1000). After washing, these were mounted in 1.2% agarose/PBS. Imaging was carried out using a Zeiss LSM 800 laser scanning confocal microscope, with a 40x water immersion objective.

### Enucleation

5 day-old fish were anaesthetized in Ringer’s saline containing buffered tricaine. The eyes were removed using electrolytically sharpened tungsten needles. Fish were allowed to recover for several hours in anesthetic-free saline. Activity was recorded 2 - 4 hours after eye removal. To enable lateral imaging of the thalamus (Fig. 5c,d), one eye was removed using this method.

### Optogenetic stimulation

5 dpf *GAL4s1020t, UAS:ChR2-eYFP, elavl3:GCaMP6f* larvae were used. All experiments were performed on fish lacking eyes. Fish were mounted in 1.2% agarose in Ringer’s saline, and imaged using two-photon microscopy as described above, at 1 Hz. Optical stimulation was carried out using a 50 µm fiber optic probe (Doric Lenses). The probe was held with a pipette holder (UT-2, Narishige), and the tip was positioned approximately 20 µm from fish, at the level of the thalamus, using a hanging drop micromanipulator (MO-202U, Narishige). The 465 nm LED (Doric) was driven with a current of 900 mA, 30 seconds after the start of imaging. 10 pulses were provided, with a pulse duration of 25 milliseconds and a frequency between 1 and 8 Hz. Each fish was exposed to at least 3 pulse trains. For Fig. 9b-c, the average of the first 29 frames was used as a reference. The ratio of all frames relative to this reference was obtained using FIJI (RRID:SCR_002285). The analysis to generate Fig. 9g was blind to the genotype.

### Laser ablation

*elavl3:GCaMP6f* larvae were anaesthetized and then mounted in 2% low-melting temperature agarose. First, the response of dorsal habenula neurons to light pulses was recorded. Lesions were then created with the femto-second laser tuned to 960 nm and fixed on a single point. Several pulses, each lasting 100 - 500 msec, were used. Lesioning was monitored by time-lapse imaging GCaMP6f fluorescence before and after each pulse, and was terminated when a cavitation bubble was seen; this was visible by simultaneously collecting light at 595 nm. Animals with bleeding in the brain after lesioning, due to bursting of a blood vessel in the thalamus, were discarded. The dorsal habenula was then re-imaged at the focal plane that was initially recorded, as determined by the focus motor, with care taken to ensure that cell shapes matched.

### Vertical migration

As described elsewhere [32], six naive larvae were tested simultaneously. Fish were placed individually in a chamber (3 cm L x 1 cm W x 5 cm H). After 3 min of adaptation to light and habituation to the chamber, 6 cycles of alternating light/dark were delivered, each consisting of 1 min light ON and 1 min light OFF. A green (24V, 525 nm peak, TMS-lite) or blue (470 nm peak, TMS-lite) LED backlight was the only visible light source in the incubator. The intensity of the green light was 3.8 mW/cm^2^, while the intensity of blue light was 6.0 mW/cm^2^, as measured using a Thorlabs light meter (PM100A and S120VC). Videos were taken at 17 fps, 1096 x 1096 pixel resolutions, using custom-written Python codes for real-time tracking of the fish position in the tank. The codes also control a USB3.0 Basler camera (acA2040-90umNIR) attached with a 1:1.8/4 mm lens (Basler) and a 830 nm longpass filter (MIDOPT, LP830) for capturing images at the IR range. Four infrared LED bars (850 nm peak TMS-lite) were used for illumination. The LED backlight was controlled by Python codes driving a microcontroller board (Arduino Uno) connected to a power supply switch (TMS-lite). The entire experiment for one transgenic line was carried out in one afternoon (3-6 pm). A total of 57 fish were tested (18 GtACR1, 17 control siblings, tested at 8 dpf; 10 GtACR2, 12 control siblings, tested at 11 dpf).

Expression of GtACR1 or GtACR2 in each fish was determined after the experiment using a fluorescent stereo microscope. No fish was excluded from analysis. The x-y coordinate data were analyzed using custom-written macros in Excel (Microsoft). The correlation coefficient of each fish (Fig.10c, e) was calculated using the *correl* function in Excel to correlate the vertical position of the fish in the tank (normalized from 0-bottom to 1-top) with the LED backlight status (0-OFF and 1-ON). To determine the initial movement of the fish upon each light offset especially when the fish was in the middle of the tank (defined as between 0.25 and 0.75 in the y-axis), we calculated the position of the fish at the 6th sec after light offset (i.e after the first 10% of darkness). Upward movement is defined as vertical position at t_6_ > t_0_ (red dots in Fig.10d, f) and downward movement is defined as vertical position at t_6_ < t_0_ (blue dots in Fig.10d, f). Because each fish has more than 1 data point in the 6 ON/OFF cycles in Fig.10d, a multilevel analysis was conducted to rule out the nested cluster (fish). Locomotion was calculated as distance moved by each fish under light ON and OFF and averaged across 6 cycles.

## Declarations

### Availability of data and material

The code and data used are available in the Figshare repository, https://figshare.com/s/68c165f8868eca15cbb9.

### Author contributions

Experiments were designed by CRK, QL and SJ. CRK carried out two-photon imaging, and optogenetic experiments with anion channelrhodopsins. SK developed software and analyzed imaging data. QL performed parapineal lesion. CK generated the *UAS:GCaMP6s* line. SJ performed wide-field imaging and analysis, dye tracing, antibody label, ChR2 manipulation and wrote the manuscript.

## Acknowledgements

We thank David Hildebrand and Isaac Bianco for providing the *elavl3:GCaMP6f* line, and Claire Wyart for providing the *UAS:ChR2-eYFP* line. The drawing in Fig. 5b was obtained from www.uoneuro.uoregon.edu.

## Figure legends

**Fig. 1 Overview of the habenula response to onset and offset of light**

**a, b** Dorsal view of the fore and midbrain of 5 day old *elavl3:CaMP6f* fish, imaged with widefield fluorescence microscopy at 200 Hz. The time since start of illumination is shown at the top right. The wedge indicates the ratio of fluorescence relative to the first frame. There is an increase in GCaMP6f fluorescence in the left habenula (arrowhead in b). **c**. Maximum ΔF/F_0_ value in the left and right habenula after onset of light in five different fish. Each circle is one fish and the line joins data points from the same fish. **d-f** Two photon imaging of the habenula in a 7dpf *GAL4s1011t, UAS:GCaMP6s* fish, at 13 Hz. **d** Average of the time-lapse sequence, showing anatomy. The dorsal left neuropil is indicated by the yellow arrowhead. **e** Spatial distribution of responses to pulses of light. Pixels are color-coded according to the temporal pattern of response, as indicated in panel f. **f** Centers of clusters obtained from running *K-means* on the time series of pixels in panel d. Cluster centers are colored in shades of blue for responses to light onset (On) and magenta and orange for responses to light offset (Off). The presence of light is indicated by the blue bars. **g, h** Neuropil response summarized from imaging 10 fish exposed to 7 pulses of blue light. **g** Pixels in the left neuropil from all fish could be classified into three main classes. They are pseudocolored and overlaid on an average image of a 6 dpf fish. The largest response was a transient response to light on (blue). A sustained response to light On (cyan) and Off (magenta) were also seen. Responses were reproducible in all ten fish. **h** Average traces obtained from neuropil pixels, shown here for two pulses of light. **i** Percentage of cells active to light On and Off in the habenula is correlated with the percentage of active pixels in neuropil. The transient and sustained neuropil responses were combined into ON. Percentage of active cells or pixels were calculated by dividing the number of cells/pixels active to the stimulus by the total number of segmented cells or neuropil pixels. Each circle per category is one fish. The bold lines show best fit (linear regression). r is the correlation coefficient. **j-k** Cumulative probability of peak ΔF/F_0_ response in cells and neuropil pixels responding to light On (j) and light Off (k). The response in the neuropil precedes the response in cells. P-value and test statistic (D) were obtained by a KS test between categories in the first 5 seconds. In panel j, the dark gray * (left) is the result of comparison between neuropil transient On and cell On, while light gray * (right) between neuropil sustained On and cell On. rHb: right habenula; lHb: left habenula. Pa: pallium; OT: optic tectum. a: anterior; p: posterior. Scale bar = 25 µm.

**Fig. 2 Response of habenula neurons to pulses of light**

**a-d** The dorsal habenula response to 7 pulses of blue light in n=10 fish *(GAL4s1011t, UAS:GCaMP6s*, 5-7 dpf). **a** Heatmaps from 5 example fish showing responses in cells that were classified as On, Off or Inhibitory (Inh). The colors indicate ΔF/F_0_, as shown in the colorbar. Responses in each fish are sorted in ascending order of mean ΔF/F_0_. Black horizontal lines separate each fish. The bold vertical lines correspond to light onset while the dashed lines indicate offset. The presence of light is also indicated by the blue bars. The height of the heatmaps represent the number of cells as indicated by the vertical scale bar. **b** Overlay of cells segmented from all fish. A small circle was drawn around the centroid of the segmented cell. Three main classes of activity are shown. Blue indicates cells responding to light ON (ON), green cells are inhibited by light (Inh) and magenta cells are activated in the absence of light (OFF). Hollow circles did not show an evoked response. The gap in the left habenula indicates the neuropil region. **c** Averaged traces from the cells in panel b, showing the response of different classes for the first two pulses of light. **d** Boxplots showing distribution of cells responding to different classes in the left and right habenula. Each circle is one fish and the line joins data points from the same fish. **e** K-means clustering of pixels in the habenula of fish from 5-10 dpf as indicated. Pixels are colored blue if they respond to light ON and magenta if they respond to light OFF. Data from each fish was analysed separately. Individual traces of the cluster centroids are not shown here but are similar to Fig. 1f. All fish have a response to light onset in the dorsal left neuropil.

**Fig. 3 Anatomical characterization of thalamic neurons projecting to the habenula**

**a** An example of DiD injection (cyan) into the dorsal neuropil of the left habenula (yellow arrowhead). The dorsal neuropil of the right habenula contains afferents from the entopeduncular nucleus (labelled by the SqKR11Et line; magenta) and has no DiD labelled neurons, indicating specificity of the injection. **b** Dorsal view of the thalamic region of a 7 day old fish following DiD injection into the dorsal left habenula neuropil. Arrowheads indicate retrogradely DiD-labelled neurons that express eGFP (shown in yellow) under the control of the vGlut2a GAL4 driver. **c** Lateral view of the fish in panel b, showing DiD labelled neurons on the right side of the brain. **d** Dorsal view of another larva, in which the dorsal right habenula had been injected with DiD. Retrogradely labelled neurons are located in the entopeduncular nucleus. **e** A double transgenic fish, with glutamatergic neurons shown in green, and GABAergic neurons shown in magenta. The neuropil in the anterior thalamus (arrow) contains magenta label (e’’), indicating the presence of GABAergic fibers. **f** Lateral view of a 7 dpf larva following injection of CM-DiI into the dorsal left habenula and labelling with anti GAD 65/67. The region of the neuropil containing CM-DiI labeled neurites (red; arrowheads; f’’) is labelled with the GAD65/67 antibody (cyan; f’). All panels except **c** are single optical sections. Pa: pallium; rHb: right habenula; lHb: left habenula; fr: fasciculus retroflexus; Th: thalamus. EN: entopeduncular nucleus; OT: optic tectum; otr: optic tract; ZLI: zona limitans intrathalamica. Scale bar = 25 µm. a: anterior; p: posterior; d: dorsal; v: ventral.

**Fig. 4 The response of thalamic neurons to irradiance change.**

**a-e** Activity in five different focal planes of a 5-day-old fish expressing GCaMP6s in thalamic neurons (arrows). Numbers indicate depth. The colours represent K-means cluster centers shown in panel **f**, with blue indicating ON responses and magenta indicating OFF responses; cyan pixels have a response to ON and OFF. **g-j** Quantitative analysis of response of anterior thalamic neurons of *GAL4s1020t, UAS:GCaMP6s* fish (5-6 dpf) to pulses of blue light. Note that this driver is not expressed in afferent retinal ganglion cells. **g.** Responses in cells from 10 fish at three different focal planes. Four pulses of blue light were given and imaging was done at 7 Hz. **g.** Heatmaps of individual cells in five example fish, showing major classes of responses seen in thalamic neurons: excitation to light ON, light OFF, or both ON and OFF (yellow). g and g’ have different scales. **h.** Segmented cells in all fish overlaid and colored by their response. **i.** Traces showing mean responses of cells in panel **h** for two blue pulses. **j** Percentage of cells responding to light ON, OFF or both ON and OFF. **k.** Neuropil responses to pulses of light. Pixels with different response classes from all fish were pseudo-colored and overlaid on an average image from a 5dpf, *GAL4s1020t, UAS:GCaMP6s* fish. **l** Average traces of responses in panel k. **m.** Percentage of neuropil pixels responding to light ON, OFF or to both ON and OFF. **n-r** Thalamus response to blue and red light. **n.** Spatial distribution of responses, color coded according to the *k-means* cluster centers in panel **o**, with blue pixels showing a sustained response to light ON, while magenta pixels and orange pixels are a mixture of responses to both ON and OFF. Z is the Z-score. **p.** Heatmaps of cells responding to 3 pulses of blue light followed by 3 pulses of red light in n=6 *GAL4s1020t, UAS:GCaMP6s* fish. Cells were classified as responding to light ON or OFF. While the same cells responded to both blue and red light, the amplitude of responses were lower to red light. **q.** Peak amplitude of response during light ON and OFF is higher to blue light than red light. Each circle represents one fish and lines join data points from the same fish. Crosses and diamonds represent median amplitude. **r.** Histogram showing amplitude of responses during blue (blue traces) and red (red traces) light ON (left panel) and OFF (right panel). Each trace is response distribution from all cells in a single fish. P-values and test statistic (D) were obtained using KS-Test on cumulative response distribution from all fish shown in the inset in r. Th:Thalamus.

**Fig. 5 Anatomical and physiological characterization of the anterior thalamic neuropil**

**a.** Lateral view of a 6 day old fish following injection of DiD (cyan) into the dorsal neuropil of the left habenula and DiI (yellow) into the right retina. Arrows indicate terminals from retinal ganglion cells in the vicinity of fibers from habenula afferents. See Movie 5. **b** Illustration of a fish larvae, showing the region imaged in panel a (red box) and in panels c and d (black box). **c-h** Response in the anterior thalamic neuropil to pulses of light. **c** Average projection of a lateral view of an *elavl3:GCaMP6f* fish, showing the thalamic neuropil (arrowhead). **d, e** The response to four pulses of blue light. Colours show the *K-means* cluster centers represented in panel e. The regions responding to light ON and light OFF are distinct in the thalamic neuropil. Responses can also be seen in the habenula. **f-h** Quantitation of the anterior thalamus neuropil response to light pulses in 8 fish. **f** Contours show a bivariate kernel density estimate of neuropil pixel location for responses to ON (shades of blue) and OFF (shades of red) of blue light in n=8 fish. The two variables here are x and y of neuropil pixels. The orientation is same as panel d. Crosses indicate median location of response to light ON, while diamonds indicate median location of response to light OFF in each fish. The dorso-ventral and anterior-posterior positions of the median centers are shown in panels **g** and **h** respectively. Each circle is one fish and lines join data points from the same fish. These panels show that ON and OFF responses have a different location, with OFF responses in a more anterior-ventral location. Scale bar = 25 µm. P-values and test statistic (D) were obtained using KS-Test on cumulative distribution of pixel location to light ON and OFF from all fish. a: anterior; p: posterior; d: dorsal; v: ventral. Hb: Habenula.

**Fig. 6 Effect of eye removal on thalamus response to light ON and OFF**

**a-i** Activity in the thalamus in response to pulses of blue light in control (n = 3) and enucleated (n = 4) fish. **a,b** Heatmaps showing activity in individual cells in control and enucleated fish classified as having response to light ON or OFF. **c** A comparison of the percentage of active cells in control and enucleated fish. The response to light ON is reduced in fish lacking eyes, while the response to light OFF is comparable to controls. **d-e.** Histogram of mean response amplitude in cells in control and enucleated fish during (d) light ON and (e) OFF. Each trace is one fish. The amplitude of response to light ON is reduced in enucleated fish. Insets show cumulative histogram from all fish. **f-i** Pixels in the anterior thalamic neuropil of control (f) and enucleated (h) fish, that are active to light ON (cyan) or OFF (pink), were combined and overlaid. Panels f and h show a dorsal view of the thalamus. The average traces from the colored pixels in f and h are shown in g and i respectively. Control fish have a response to light ON and OFF, whereas enucleated fish only have a response to light OFF. **j** Percentage of active neuropil pixels in control and enucleated fish. **k-l** Cumulative probability of mean response amplitude in pixels of control and enucleated animals to light ON (k) and light OFF (l). Mean response during light OFF is not significantly different in enucleated and control fish. **m** Dorsal view of a 6 day old fish, following injection of DiD into the dorsal left habenula neuropil and CM-DiI into the pineal organ. See movie 6 for the complete z-stack. **n** Lateral view of the right side of a 6 day old fish, showing anterogradely labeled fibers from the pineal (red) and retrogradely labeled fibers from the habenula (cyan). The arrow indicates a pineal axon in the neuropil of the anterior thalamus. **o, p** Response in the pineal organ to pulses of blue light (n=4 fish). Only OFF responses can be detected. (o) Pixels showing OFF responses are combined from all fish and overlaid. (p) Average trace from the colored pixels. The habenula is shown here for orientation only; habenula neuron responses have been masked. **q-s** Responses in the habenula to light OFF in control (q) and enucleated (r) fish (*GAL4s1011t, UAS:GCaMP6s*, n=4 fish). Each row in the heat maps represents an individual cell. **s** Percentage of cells showing an OFF response. Each circle is one fish and lines join data points from same fish before and after enucleation. Although reduced in number, there are still cells that display an OFF response. D-statistic and p-values in panels d, e, k and l were obtained using the KS test on response amplitude distribution. Panels f, h, m and o are single optical sections; n is a projection spanning 19.25 µm. rHb:right habenula; lHb: left habenula; Th: thalamus; a: anterior; p: posterior; d: dorsal; v: ventral. Scale bar = 25 µm.

**Fig. 7 The effect of lesioning the anterior thalamic neuropil on habenula response to light.**

**a** Dorsal view of an *elavl3:GCaMP6f* fish, showing lesion bubbles in the anterior thalamic neuropil created by a femtosecond laser (arrows). The bubble reflects the two photon laser, and is thus captured in a separate channel from GCaMP6f fluorescence. **a’,a’’** Close-up of the anterior thalamus neuropil before (a’) and during (a’’) lesion. The cavity has not yet formed. **b** Heat map showing habenula cell responses before (left) and after (right) lesioning in 3 fish. **c, d** The cells segmented from all 3 fish are drawn as circles and overlaid. Responding cells before and after lesion are colored as indicated. **e, f** Histogram showing distribution of mean intensity in habenula neurons during light ON (e) and OFF (f) before and after lesion. Insets show cumulative distribution from all fish. P-values and test statistic were obtained using the KS-test. **g** Comparison of percentage of cells responding to light ON and OFF before and after lesion. Each circle is one fish and lines join data points from the same fish. a: anterior; p: posterior; Images are all single optical sections. Scale bar = 25 µm.

**Fig. 8 GtACR1 expression in the thalamus disrupts habenula response to light ON and OFF**

**a** A 6 day old fish expressing GtACR1-eYFP under the control of the *GAL4s1020t* driver. GtACR1-expressing cells in the anterior thalamus (colored orange-purple) are indicated by the yellow arrowheads. Puncta of GtACR1-eYFP are visible. There is a low level of GCaMP6f expression, shown in green here. **b** A more dorsal focal plane, with habenula afferents labelled by the *sqKR11Et* line (magenta). GtACR1-eYFP puncta are indicated by the arrowheads. The asterisks indicate autofluorescent pigment cells. **c-k** Comparison of dorsal left habenula neuropil response to pulses of blue light, in controls and fish expressing GtACR1 in the thalamus. **c,d** Response in the neuropil of control (**c**; n = 7) and GtACR1 expressing siblings (**d**; n = 11). -Blue represents fast ON, cyan represents slow ON, whereas magenta represents OFF response. **e,f** The average of the colored pixels from c and d respectively. Shaded areas show 95% confidence intervals. **g,h** Boxplots show the distribution of percentage of neuropil pixels showing a response to light ON or OFF in fish expressing GtACR1 (h) or control siblings (g). Each circle is one fish and lines join data points from same fish. P-values and test statistic are obtained using Mann-Whitney U test between the distribution of pixels in control and GtACR1 fish of the same response class. **i,j** Histograms showing neuropil response to light ON and OFF in individual fish expressing GtACR1 (j) and control siblings (i). There is a leftward shift in the distribution for response to light ON in GtACR1 fish. **k** Interpolations of the histograms in panels i-j using a smoothing spline fit to show the overall distribution per category. Pa: pallium; Th: thalamus; hc: habenula commissure; rHb: right habenula; lHb: left habenula; a: anterior; p: posterior.

**Fig. 9 Effect of optogenetic stimulation of the thalamus on habenula activity.**

**a** Expression of ChR2-eYFP in the thalamus (arrowheads) of a 5 day old *GAL4s1020t, UAS:ChR2-eYFP, elavl3:GCaMP6f* fish. **b, c** Activity in the habenula of a ChR2-expressing fish, with (b) and without (c) blue LED stimulation of the thalamus. The images show the maximum projections of F/F_0_ images for a 25-second period after blue LED illumination, following subtraction of maximum projections of the period before illumination (i.e. difference in activity before and after stimulation). **d-f** Heatmaps showing temporal activity from habenula neurons segmented in fish with (e-f) and without (d) ChR2. In panels e (n = 3 fish) and f (n = 2 fish), blue light pulse was given at the time indicated by the black dashed line and at a frequency specified. **g.** Cumulative distribution of mean response amplitude, 10 seconds after stimulation in ChR2-expressing and control fish and a randomly chosen 10 second period in fish with no stimulation. All stimulation frequencies were combined. The fish with ChR2 show increased ΔF/F_0_ after optogenetic stimulation. Test statistic and p-values were obtained using KS-test. The gray * (bottom) is the result of comparison between control siblings and Chr2-expressing fish, while the black * (top) is between no stimulation and Chr2-expressing fish. **h** Mean amplitude before and after optogenetic stimulation at different frequencies. Each square stands for a stimulus trial. Scale bar = 25 µm. Pa: pallium, a: anterior, p:posterior, lHb: left habenula, rHb: right habenula.

**Fig. 10 Anion channelrhodopsin expression in the thalamus disrupts vertical migration to irradiance change**

**a,b** Vertical position of control and *GAL4s1020t, UAS:GtACR1-eYFP* fish exposed to alternating periods of light and darkness. Thin lines show trajectories of individual fish, while the thicker red line indicates the average. Shading indicates 95% confidence intervals. Overall, control fish move up when lights go ON and down when lights go OFF. GtACR1-expressing fish have the opposite behavior. **c,e** Correlation between light (1 for ON, 0 for OFF) and vertical movement. Error bars indicate 95% confidence interval. Correlation is high for controls, but not for GtACR1 (c) or GtACR2 (e) expressing fish. **d,f** Direction of initial vertical movement at light OFF for all fish, at all transitions. Blue indicates downward movement, while red indicates upward movement. GtACR1 (d) and GtACR2 (f) expressing fish tend to move upwards at light OFF, whereas non-expressing siblings tend to move down. **g,h** Amount of movement of GtACR1 (g) and GtACR2 (h) expressing fish at light ON and OFF averaged across all 6 cycles.

## Supplementary Figures

**Fig. S1. The effect of parapineal lesion on habenula response to blue light.**

**a** Visualization of the parapineal (yellow arrow), which is located adjacent to the left habenula and innervates the dorsal neuropil. **b, c** Two photon lesioning of the parapineal. **b** Before lesioning. **c** After lesioning, which led to formation of a bubble (arrow). **d, e** Habenula cells segmented from 5 fish, overlaid on top of each other, showing responses before and after lesion. Cells responding to light ON are shown in blue and to OFF in pink. **f, g** Heat maps of the habenula cells, in the 5 fish, responding to light ON and OFF before (f) and after (g) lesioning the parapineal. Horizontal black lines divide data from different fish. **h** Percentage of cells showing ON and OFF responses before and after parapineal lesioning. **i, j** Histogram showing distribution of mean intensity in habenula neurons during light ON (e) and OFF (f) before and after lesion. Insets show cumulative distribution from all fish. P-values and test statistic were obtained using the KS-test. pp:parapineal, lHb: Left Habenula, rHb:Right Habenula, cv: circumventricular organ; a: anterior, p:posterior. Scale bar = 25 µm.

**Figure S2. Examples of signals that were excluded from visualisation of k-means clusters.**

**a-e** Pixels showing stimulus-independent activity in the thalamus, at 5 different focal planes. Pixels are colored according to the traces in panel **f**. For clarity, these signals were excluded from the visualisation of clusters representing light evoked activity shown in Fig. 4a-e. **g** Stimulus-independent activity in the habenula. Pixels are colored according to the traces in panel **h.** For clarity, these signals were excluded from the visualisation of clusters representing light evoked activity shown in Fig. 1e-f. **f**, **h** Cluster centers that did not represent light-evoked activity in the habenula, obtained by running *K-means* on the time series of pixels in panel a-e and g. Th: Thalamus, a:anterior, p:posterior. Scale bar = 25 µm.

## Movie legend

**Movie 1. Z-stack through a brain following DiD injection into the dorsal left habenula neuropil**.

DiD label (cyan) is seen bilaterally in two clusters of neurons in the anterior thalamus, starting from a depth of 65 µm from the first plane. Sparse labeling can also be seen in the ipsilateral entopeduncular nucleus (EN), at a depth of about 100 – 110 µm. Glutamatergic neurons are labeled by *vGlut2a:GAL4,UAS:eGFP* (yellow), while GABAergic neurons are labeled by *gad1b:DsRed* (magenta). The left fasciculus retroflexus is labeled by axons from the habenula. This is a dorsal view, with anterior to the left. Gamma = 0.45.

**Movie 2. Z-stack through the brain following DiD injection into the dorsal right habenula neuropil**

Retrogradely labeled cells are seen primarily in the ipsilateral entopeduncular nucleus (arrow). Labelled axons are also visible in the neuropils of the left habenula. These may arise from neurons that innervate the anterior right thalamus and/or from the right entopeduncular nucleus (arrow). This is a dorsal view, with anterior to the left.

**Movie 3. Z-stack of 6 day old gad1b:DsRed, vglut2a:GAL4, UAS:eGFP fish**. GABAergic neurons (magenta) are visible in the thalamus, below the habenula. Arrows indicate the anterior thalamic neuropil which contains DsRed-labelled fibers (~50 µm below the first plane). The entopeduncular nucleus does not contain DsRed-labelled fibers. In the first frame, DsRed-labelled neurites are visible in the optic tectum, but not in the habenula neuropil. Anterior is to the left. The stack goes from dorsal to ventral. rHb: right habenula; lHb: left habenula; OT: optic tectum; EN: entopeduncular nucleus.

**Movie 4. Lateral view of a fish following DiD injection in the dorsal left neuropil of the habenula**. The right side of the fish shown in Movie 1. Thalamic neurons that have been retrogradely labeled are shown in cyan. Glutamatergic neurons are labeled by *vGlut2a:GAL4,UAS:eGFP* (yellow), while GABAergic neurons are labeled by *gad1b:DsRed* (magenta). DiD labeled cells extend neurites into neuropil of the anterior thalamus. A number are labeled by eGFP (arrows), but none are labeled by DsRed. The optic tract is visible in the DIC image, and contains eGFP labeled axons. Note that anterior is to the right in this stack.

**Movie 5. Lateral view of the left anterior thalamus following injection of DiD (cyan) into the dorsal left habenula neuropil and DiI (yellow) into the right eye**.

The arrow shows intermingling of retinal and habenula afferent fibers in the thalamic neuropil. The stack runs from lateral to medial, and habenula afferents and RGC terminals meet anteriorly and medially to the optic tract. fr: fasciculus retroflexus. This is a 6-day-old fish, with anterior to the left.

**Movie 6. Z-stack of a 6 day old fish following CM-DiI injection into the pineal and DiD into the dorsal left habenula**.

Pineal axons (red) project laterally and then posteriorly. Arrows indicate axons that enter anterior thalamic neuropil, where retrogradely labeled fibers from the habenula (cyan) are visible. This is a dorsal view, with anterior to the left.

## REFERENCES

1 Stephenson-Jones M, Floros O, Robertson B, Grillner S. Evolutionary conservation of the habenular nuclei and their circuitry controlling the dopamine and 5-hydroxytryptophan (5-HT) systems. Proc. Natl. Acad. Sci. U. S. A. 2012;109:E164–73.

2 Lee A, Mathuru AS, Teh C, Kibat C, Korzh V, Penney TB, et al. The habenula prevents helpless behavior in larval zebrafish. Curr. Biol. 2010;20:2211–6.

3 Agetsuma M, Aizawa H, Aoki T, Nakayama R, Takahoko M, Goto M, et al. The habenula is crucial for experience-dependent modification of fear responses in zebrafish. Nat. Neurosci. 2010;13:1354–6.

4 Zhang J, Tan L, Ren Y, Liang J, Lin R, Feng Q, et al. Presynaptic Excitation via GABAB Receptors in Habenula Cholinergic Neurons Regulates Fear Memory Expression. Cell. 2016;166:716–28.

5 Matsumoto M, Hikosaka O. Lateral habenula as a source of negative reward signals in dopamine neurons. Nature. 2007;447:1111–5.

6 Matsumoto M, Hikosaka O. Representation of negative motivational value in the primate lateral habenula. Nat. Neurosci. 2009;12:77–84.

7 Amo R, Fredes F, Kinoshita M, Aoki R, Aizawa H, Agetsuma M, et al. The habenulo-raphe serotonergic circuit encodes an aversive expectation value essential for adaptive active avoidance of danger. Neuron. 2014;84:1034–48.

8 Fowler CD, Lu Q, Johnson PM, Marks MJ, Kenny PJ. Habenular α5 nicotinic receptor subunit signalling controls nicotine intake. Nature. 2011;471:597–601.

9 Aizawa H, Cui W, Tanaka K, Okamoto H. Hyperactivation of the habenula as a link between depression and sleep disturbance. Front. Hum. Neurosci. 2013;7:826.

10 Chou M-Y, Amo R, Kinoshita M, Cherng B-W, Shimazaki H, Agetsuma M, et al. Social conflict resolution regulated by two dorsal habenular subregions in zebrafish. Science. 2016;352:87–90.

11 Golden SA, Heshmati M, Flanigan M, Christoffel DJ, Guise K, Pfau ML, et al. Basal forebrain projections to the lateral habenula modulate aggression reward. Nature. 2016;534:688–92.

12 Jhou TC, Fields HL, Baxter MG, Saper CB, Holland PC. The rostromedial tegmental nucleus (RMTg), a GABAergic afferent to midbrain dopamine neurons, encodes aversive stimuli and inhibits motor responses. Neuron. 2009;61:786–800.

13 Quina LA, Tempest L, Ng L, Harris JA, Ferguson S, Jhou TC, et al. Efferent pathways of the mouse lateral habenula. J. Comp. Neurol. 2015;523:32–60.

14 Morley BJ, Spangler KM, Javel E. The development of somatostatin immunoreactivity in the interpeduncular nucleus of the cat. Brain Res. 1985;352:241–8.

15 Wang RY, Aghajanian GK. Physiological evidence for habenula as major link between forebrain and midbrain raphe. Science. 1977;197:89–91.

16 Appelbaum L, Wang GX, Maro GS, Mori R, Tovin A, Marin W, et al. Sleep-wake regulation and hypocretin-melatonin interaction in zebrafish. Proc. Natl. Acad. Sci. U. S. A. 2009;106:21942–7.

17 Hong S, Hikosaka O. The globus pallidus sends reward-related signals to the lateral habenula. Neuron. 2008;60:720–9.

18 Dreosti E, Vendrell Llopis N, Carl M, Yaksi E, Wilson SW. Left-right asymmetry is required for the habenulae to respond to both visual and olfactory stimuli. Curr. Biol. 2014;24:440–5.

19 Krishnan S, Mathuru AS, Kibat C, Rahman M, Lupton CE, Stewart J, et al. The right dorsal habenula limits attraction to an odor in zebrafish. Curr. Biol. 2014;24:1167–75.

20 Miyasaka N, Morimoto K, Tsubokawa T, Higashijima S-I, Okamoto H, Yoshihara Y. From the olfactory bulb to higher brain centers: genetic visualization of secondary olfactory pathways in zebrafish. J. Neurosci. 2009;29:4756–67.

21 Zhao H, Rusak B. Circadian firing-rate rhythms and light responses of rat habenular nucleus neurons in vivo and in vitro. Neuroscience. 2005;132:519–28.

22 Sakhi K, Wegner S, Belle MDC, Howarth M, Delagrange P, Brown TM, et al. Intrinsic and extrinsic cues regulate the daily profile of mouse lateral habenula neuronal activity. J. Physiol. 2014;592:5025–45.

23 Semm P, Demaine C. Electrophysiology of the pigeon’s habenular nuclei: evidence for pineal connections and input from the visual system. Brain Res. Bull. 1984;12:115–21.

24 Bianco IH, Engert F. Visuomotor transformations underlying hunting behavior in zebrafish. Curr. Biol. 2015;25:831–46.

25 Amo R, Aizawa H, Takahoko M, Kobayashi M, Takahashi R, Aoki T, et al. Identification of the zebrafish ventral habenula as a homolog of the mammalian lateral habenula. J. Neurosci. 2010;30:1566–74.

26 Burrill JD, Easter SS Jr. Development of the retinofugal projections in the embryonic and larval zebrafish (Brachydanio rerio). J. Comp. Neurol. 1994;346:583–600.

27 Robles E, Laurell E, Baier H. The retinal projectome reveals brain-area-specific visual representations generated by ganglion cell diversity. Curr. Biol. 2014;24:2085–96.

28 Turner KJ, Hawkins TA, Yáñez J, Anadón R, Wilson SW, Folgueira M. Afferent Connectivity of the Zebrafish Habenulae. Front. Neural Circuits. 2016;10:30.

29 Butler AB, Saidel WM. Clustered phylogenetic distribution of nucleus rostrolateralis among ray-finned fishes. Brain Behav. Evol. 2003;62:152–67.

30 Saidel WM. Nucleus rostrolateralis: an expansion of the epithalamus in some actinopterygii. Anat. Rec. 2013;296:1594–602.

31 Hendricks M, Jesuthasan S. Asymmetric innervation of the habenula in zebrafish. J. Comp. Neurol. 2007;502:611–9.

32 Lin Q, Jesuthasan SJ. Masking of a circadian behavior in larval zebrafish involves the thalamo-habenula pathway. Scientific Reports. 2017;7:4104.

33 Chen T-W, Wardill TJ, Sun Y, Pulver SR, Renninger SL, Baohan A, et al. Ultrasensitive fluorescent proteins for imaging neuronal activity. Nature. 2013;499:295–300.

34 Mueller T. What is the Thalamus in Zebrafish? Front. Neurosci. 2012;6:64.

35 Sherman SM, Guillery RW. The role of the thalamus in the flow of information to the cortex. Philos. Trans. R. Soc. Lond. B Biol. Sci. 2002;357:1695–708.

36 Satou C, Kimura Y, Hirata H, Suster ML, Kawakami K, Higashijima S-I. Transgenic tools to characterize neuronal properties of discrete populations of zebrafish neurons. Development. 2013;140:3927–31.

37 Moore RY, Speh JC. GABA is the principal neurotransmitter of the circadian system. Neurosci. Lett. 1993;150:112–6.

38 Venugopalan V, Guerra A 3rd, Nahen K, Vogel A. Role of laser-induced plasma formation in pulsed cellular microsurgery and micromanipulation. Phys. Rev. Lett. 2002;88:078103.

39 Vogel A, Noack J, Hüttman G, Paltauf G. Mechanisms of femtosecond laser nanosurgery of cells and tissues. Appl. Phys. B. 2005;81:1015–47.

40 Govorunova EG, Sineshchekov OA, Janz R, Liu X, Spudich JL. NEUROSCIENCE. Natural light-gated anion channels: A family of microbial rhodopsins for advanced optogenetics. Science. 2015;349:647–50.

41 Mohamed GA, Cheng RK, Ho J, Krishnan S, Mohammad F, Claridge-Chang A, Jesuthasan S. Optical inhibition of zebrafish behavior with anion channelrhodopsins. bioRxiv. 2017; doi:10.1101/158899

42 Takeuchi T, Duszkiewicz AJ, Sonneborn A, Spooner PA, Yamasaki M, Watanabe M, et al. Locus coeruleus and dopaminergic consolidation of everyday memory. Nature. 2016;537:357–62.

43 Raimondo JV, Kay L, Ellender TJ, Akerman CJ. Optogenetic silencing strategies differ in their effects on inhibitory synaptic transmission. Nat. Neurosci. 2012;15:1102–4.

44 Fernandes AM, Fero K, Arrenberg AB, Bergeron SA, Driever W, Burgess HA. Deep brain photoreceptors control light-seeking behavior in zebrafish larvae. Curr. Biol. 2012;22:2042–7.

45 Aarts E, Verhage M, Veenvliet JV, Dolan CV, van der Sluis S. A solution to dependency: using multilevel analysis to accommodate nested data. Nat. Neurosci. 2014;17:491–6.

46 Warp E, Agarwal G, Wyart C, Friedmann D, Oldfield CS, Conner A, et al. Emergence of patterned activity in the developing zebrafish spinal cord. Curr. Biol. 2012;22:93–102.

47 Yáñez J, Busch J, Anadón R, Meissl H. Pineal projections in the zebrafish (Danio rerio): overlap with retinal and cerebellar projections. Neuroscience. 2009;164:1712–20.

48 Jetti SK, Vendrell-Llopis N, Yaksi E. Spontaneous activity governs olfactory representations in spatially organized habenular microcircuits. Curr. Biol. 2014;24:434–9.

49 Steenbergen PJ, Richardson MK, Champagne DL. Patterns of avoidance behaviours in the light/dark preference test in young juvenile zebrafish: a pharmacological study. Behav. Brain Res. 2011;222:15–25.

50 Saidel WM, Butler AB. Retinal projections in the freshwater butterfly fish, Pantodon buchholzi (Osteoglossoidei). II. Differential projections of the dorsal and ventral hemiretinas. Brain Behav. Evol. 1991;38:154–68.

51 Zhang B-B, Yao Y-Y, Zhang H-F, Kawakami K, Du J-L. Left Habenula Mediates Light-Preference Behavior in Zebrafish via an Asymmetrical Visual Pathway. Neuron. 2017;93:914–28.e4.

52 Mueller T, Guo S. The distribution of GAD67-mRNA in the adult zebrafish (teleost) forebrain reveals a prosomeric pattern and suggests previously unidentified homologies to tetrapods. J. Comp. Neurol. 2009;516:553–68.

53 Hattar S, Kumar M, Park A, Tong P, Tung J, Yau K-W, et al. Central projections of melanopsin-expressing retinal ganglion cells in the mouse. J. Comp. Neurol. 2006;497:326–49.

54 Morin LP, Studholme KM. Retinofugal projections in the mouse. J. Comp. Neurol. 2014;522:3733–53.

55 Mohamed GA, Cheng R-K, Ho J, Krishnan S, Mohammad F, Claridge-Chang A, et al. Optical inhibition of zebrafish behavior with anion channelrhodopsins [Internet]. bioRxiv. 2017. Available from: http://dx.doi.org/10.1101/158899

56 Díaz C, Puelles L. Afferent connections of the habenular complex in the lizard Gallotia galloti. Brain Behav. Evol. 1992;39:312–24.

57 Kemali M, Guglielmotti V, Gioffré D. Neuroanatomical identification of the frog habenular connections using peroxidase (HRP). Exp. Brain Res. 1980;38:341–7.

58 Marburg O. The structure and fiber connections of the human habenula. J. Comp. Neurol. 1944;80:211–33.

59 Cragg BG. The connections of the habenula in the rabbit. Exp. Neurol. 1961;3:388–409.

60 Moore RY, Weis R, Moga MM. Efferent projections of the intergeniculate leaflet and the ventral lateral geniculate nucleus in the rat. J. Comp. Neurol. 2000;420:398–418.

61 Oh SW, Harris JA, Ng L, Winslow B, Cain N, Mihalas S, et al. A mesoscale connectome of the mouse brain. Nature. 2014;508:207–14.

62 Ely BA, Xu J, Goodman WK, Lapidus KA, Gabbay V, Stern ER. Resting-state functional connectivity of the human habenula in healthy individuals: Associations with subclinical depression. Hum. Brain Mapp. 2016;37:2369–84.

63 Torrisi S, Nord CL, Balderston NL, Roiser JP, Grillon C, Ernst M. Resting state connectivity of the human habenula at ultra-high field. Neuroimage. 2017;147:872–9.

64 Teh C, Chudakov DM, Poon K-L, Mamedov IZ, Sek J-Y, Shidlovsky K, et al. Optogenetic in vivo cell manipulation in KillerRed-expressing zebrafish transgenics. BMC Dev. Biol. 2010;10:110.

65 Baier H, Scott EK. Genetic and optical targeting of neural circuits and behavior–zebrafish in the spotlight. Curr. Opin. Neurobiol. 2009;19:553–60.

66 Arrenberg AB, Del Bene F, Baier H. Optical control of zebrafish behavior with halorhodopsin. Proc. Natl. Acad. Sci. U. S. A. 2009;106:17968–73.

67 Freeman J, Vladimirov N, Kawashima T, Mu Y, Sofroniew NJ, Bennett DV, et al. Mapping brain activity at scale with cluster computing. Nat. Methods. 2014;11:941–50.

68 Wyart C, Del Bene F, Warp E, Scott EK, Trauner D, Baier H, et al. Optogenetic dissection of a behavioural module in the vertebrate spinal cord. Nature. 2009;461:407–10.

